# Transcriptomic and functional genetic evidence for distinct ecophysiological responses across complex life cycle stages

**DOI:** 10.1101/2022.01.16.476527

**Authors:** Philip J. Freda, Jantina Toxopeus, Edwina J. Dowle, Zainab M. Ali, Nicholas Heter, Rebekah L. Lambert-Collier, Isaiah Sower, Joseph C. Tucker, Theodore J. Morgan, Gregory J. Ragland

## Abstract

Organisms with complex life cycles demonstrate a remarkable ability to change their phenotypes across development, presumably as an evolutionary adaptation to developmentally variable environments. Developmental variation in environmentally sensitive performance, and thermal sensitivity in particular, has been well documented in holometabolous insects. For example, thermal performance in adults and juvenile stages exhibit little genetic correlation (genetic decoupling) and can evolve independently, resulting in divergent thermal responses. Yet, we understand very little about how this genetic decoupling occurs. We tested the hypothesis that genetic decoupling of thermal physiology is driven by fundamental differences in physiology between life stages, despite a potentially conserved Cellular Stress Response. We used RNAseq to compare transcript expression in response to a cold stressor in *Drosophila melanogaster* larvae and adults and used RNAi (RNA interference) to test whether knocking down nine target genes differentially affected larval and adult cold tolerance. Transcriptomic responses of whole larvae and adults during and following exposure to -5°C were largely unique both in identity of responding transcripts and in temporal dynamics. Further, we found no relationship between stage-specificity and tissue-specificity of transcripts, suggesting that the differences are not simply driven by differences in tissue composition across development. In addition, RNAi of target genes resulted in largely stage-specific and sometimes sex-specific effects on cold tolerance. The combined evidence suggests that thermal physiology is largely stage-specific at the level of gene expression, and thus natural selection may be acting on different loci during the independent thermal adaptation of different life stages.

**Summary Statement:** RNAseq and gene knockout via transgenic RNAi lines suggest that physiological responses to low temperatures are largely distinct across life stages of the fly *Drosophila melanogaster*.

## Introduction

Many organisms developing from juvenile life stages through adulthood are faced with changing environmental conditions that differ dramatically but predictably during development. These shifting conditions may include resource availability, predator/herbivore abundance, and abiotic factors such as temperature (Krebs and Loeschcke, 1995; Ragland and Kingsolver, 2008; Woods, 2013). To survive these environmental changes, organisms may also dramatically change their morphology, behavior, and physiology across development. For example, juvenile stages often specialize for feeding and growth, while adults primarily (and sometimes exclusively) disperse and mate (Kingsolver et al., 2011; McGraw and Antonovics, 1983; Moran, 1994; Schluter et al., 1991). These developmentally-variable environments and key fitness components (e.g., growth vs. reproduction) lead to shifting natural selection, which may favor different trait combinations in different life stages (Haldane, 1932; Moran, 1994). This is perhaps most apparent in organisms that metamorphose like amphibians and holometabolous insects. Their morphology has evolved independently in juvenile and adult stages that inhabit drastically different ecological niches. There are clear physiological differences across complex life cycle stages as well, in part because distinct developmental machinery underlies distinct morphologies and life history strategies across the life cycle (Arbeitman et al., 2002; Herrig et al., 2021; van Gestel et al., 2019). Such morphological and developmental decoupling supports the adaptive decoupling hypothesis, which posits that natural selection favors reduced genetic correlation across developmental stages to allow for stage-specific adaptation (Moran, 1994).

In addition to developmental differences in ‘baseline’ physiology, physiological responses to environmental perturbations may also vary across the life cycle. Many key studies have examined developmental variation in environmental responses by manipulating temperature, a nearly universal selective factor that often varies over the course of development (Bowler and Terblanche, 2008; Jensen et al., 2007; Klockmann et al., 2017). Most of these studies show that thermal responses (survival and various metrics of performance) have very low or absent genetic correlations between juvenile and adult stages of holometabolous insects (Dierks et al., 2012; Gilchrist et al., 1997; Loeschcke and Krebs, 1996; Tucić, 1979). Indeed, our recent studies show that the genetic correlation between juvenile and adult cold hardiness in the fly *Drosophila melanogaster* are not detectably higher than zero, with no evidence for pleiotropic effects of SNP (single nucleotide polymorphism) variation on thermal performance across metamorphosis (Freda et al., 2017; Freda et al., 2019).

We reason that there are two hypotheses that could explain such extreme genetic decoupling of thermal physiology across development. The first, the ‘developmentally distinct physiology’ hypothesis, posits that environmental responses may indeed be very different across life stages, mirroring the differences in developmental regulation. In this scenario different genes would contribute to environmental responses across stages, with relatively low cross-stage pleiotropy.

Though the developmentally distinct physiology hypothesis is consistent with the observed lack of genetic correlations across life stages, it would be somewhat at odds with predictions based on the conserved cellular stress response. The Cellular Stress Response, or CSR, is an apparently conserved set of changes in cell physiology in response to a variety of environmental stressors (Kültz, 2005). For example, heat shock proteins and related chaperonins are up-regulated in response to multiple stressors, including temperatures that are relatively hot or cold compared to an organism’s optimal environmental temperature (Colinet et al., 2010b; Philip and Lee, 2010; Yocum, 2001). If these heat shock responses and other elements of the CSR have a substantial role in whole-organism level environmental responses, then many elements of environmental physiological responses should be very similar across the life cycle. Some elements of environmental physiological responses are admittedly tissue specific. For example, ion homeostasis in the gut and central nervous system has a well-established role in the response to mild low temperatures in many insect species (MacMillan et al., 2015; Overgaard and MacMillan, 2017). However, such tissue-specific responses may also contribute similarly to environmental responses across life stages.

Such conserved cellular and tissue-level responses would argue for a second, ‘developmentally conserved physiology’ hypothesis, positing that thermal physiology could be largely conserved across development, with only a few stage-specific processes harboring segregating genetic variation. This explanation is less obvious, but still consistent with the observed lack of genetic correlations for environmental physiology across life stages. In this scenario there may be many processes (e.g., the CSR) that universally affect thermal physiology across development, but genetic loci that regulate these processes are highly conserved, and thus not genetically variable. Genetic correlations only assess whether *variants* at loci affect two traits (e.g., juvenile and adult performance), not whether a given locus itself affects the traits. Thus, these conserved loci would not influence measures of genetic correlations. Rather, a subset of an environmental response may be stage-specific and mediated by genetically variable loci. This scenario could also generate low genetic correlations across life stages.

To test these two hypotheses, we examined physiological responses to cold across the life cycle in *D. melanogaster*, using two approaches to compare larvae (juveniles) and adults separated by a major metamorphic transition. First, we tested whether whole transcriptome responses to low temperature exposure differ in identity of responding transcripts and/or their temporal patterns of differential expression. Transcriptome sequencing provides a broad snapshot of organismal physiology, and allowed us to assess the similarity of the environmental (temperature) response across the two life cycle stages. Second, we tested whether knocking down a set of nine candidate genes affected response to low temperature in larvae, adults, or both. We selected these candidates based on a previous study that found evidence for knockout effects on cold performance in adult *D. melanogaster* (Teets and Hahn, 2018). Though the sample of nine genes is relatively small, it provides a first functional test for the presence of stage-specific (consistent with the developmentally distinct physiology hypothesis) or cross-stage (consistent with the developmentally conserved hypothesis) genetic effects on environmental physiology regardless of genetic variability.

## Materials and Methods

### Fly rearing

We obtained all *D. melanogaster* (Meigen) lines (Table S1) from the Bloomington Drosophila Stock Center (BDSC; Bloomington IN, USA) at Indiana University – specific lines used in this study are described below. We reared flies at 25°C, 12:12 L:D in narrow vials on media containing agar, cornmeal, molasses, yeast, and antimicrobial agents propionic acid and Tegosept (Genesee Scientific, Morrisville NC, USA), as described previously (Freda et al., 2017; Freda et al., 2019). We sorted parental flies from appropriate lines (details below) under light CO_2_ anesthesia and transferred them into fresh vials containing media sprinkled with dry, active yeast to facilitate oviposition. We then transferred the parents each day for four consecutive days into fresh vials to produce offspring for use in experiments. The vials from the first egg-laying day were discarded to remove any residual effect of anesthesia on oviposition. We collected third instar larvae and 5 d-old adults for use in both experiments described below. We extracted experimental third instar feeding larvae from cultures 5 d post-oviposition using a 20% w/v sucrose solution and following the protocol of Freda et al. (2017). Experimental adults were collected and sorted into fresh vials under light CO_2_ anesthesia 10 - 12 d post-oviposition (within 1 - 2 d of eclosion). These flies were held at 25°C, 12:12 L:D until 5 d-old to limit any carryover effects of CO_2_ exposure (Nilson et al., 2006).

### Experiment 1: Whole transcriptome response to low temperature

To obtain a transcriptomic metric for how physiology changes during cold exposure and subsequent recovery under benign conditions, we sampled whole-body transcriptomes of third instar larvae and 5-day old adult *D. melanogaster* prior to, during, and after exposure to a cold temperature (Fig. 1A).

**Figure 1.**
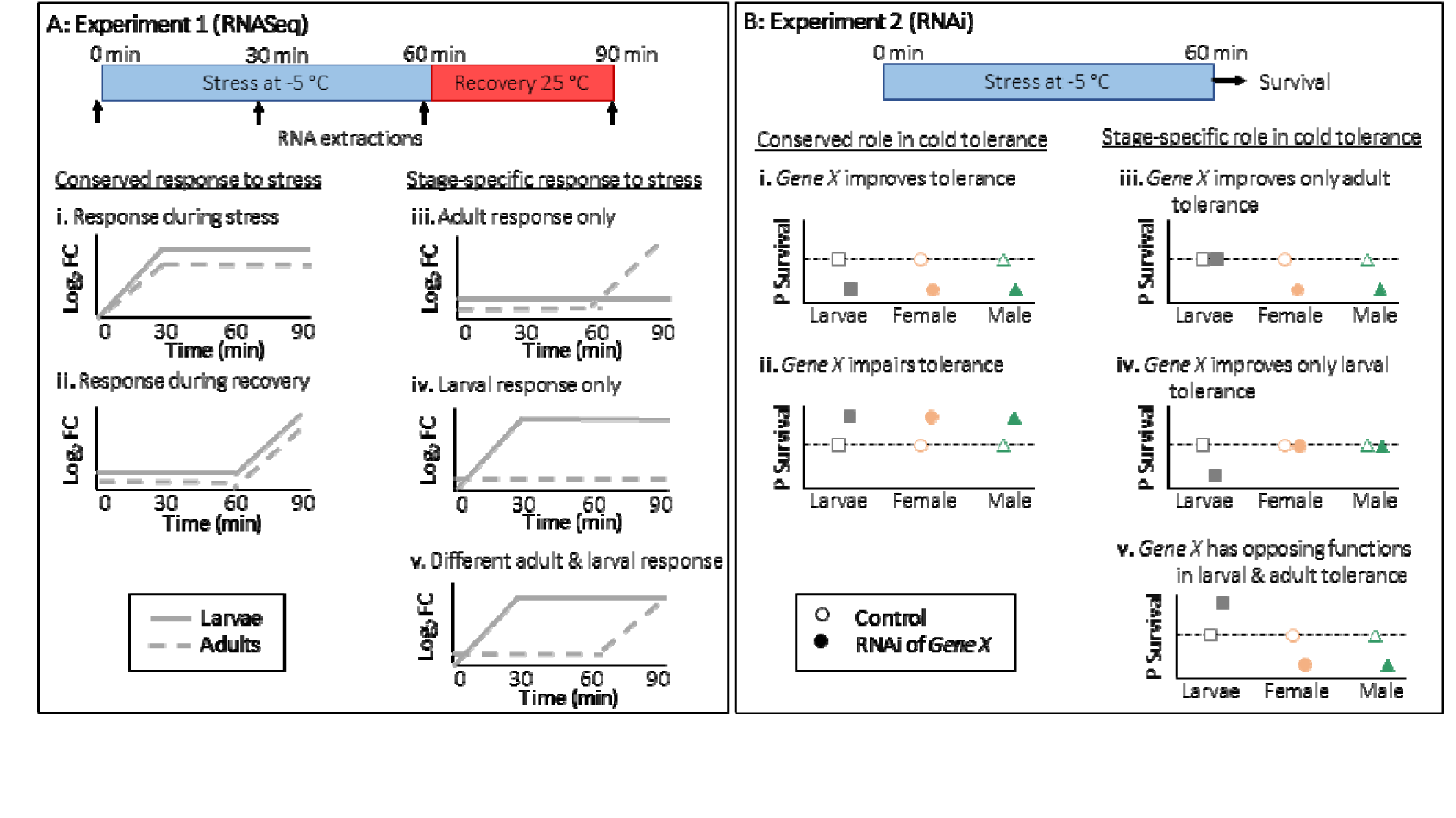
Summary of methods and example predictions for Experiments 1 and 2. Time courses at the top of each column show sampling time points during the **(A)** RNAseq experiment or the treatment prior to the measure of survival in the **(B)** RNAi line experiments. Panel **A** illustrates predicted patterns of differentially-expressed genes in *Drosophila melanogaster* during and after cold stress, with examples of (i, ii) larvae and adults exhibiting a conserved transcriptional response to cold and (iii, iv, v) larvae and adults exhibiting different transcriptional responses to cold. Predictions (log2 fold change, FC) do not differentiate between up- and down-regulated transcripts. Panel **B** illustrates predicted effects of target gene (*Gene X*) RNA interference (RNAi) on the proportion of surviving *D. melanogaster* after cold stress compared to control flies (no RNAi), with examples of the RNAi having a similar effect on cold tolerance of larvae, female adults, and male adults (i, ii) and the RNAi having life stage-specific effects on cold tolerance (iii, iv, v). We predict that knocking out a gene with a positive effect (improves cold tolerance) decreases survival of RNAi lines, whereas knocking out a gene with a negative effect (impairs tolerance) increases survival of RNAi lines.

We crossed five male and five virgin female flies from each of six Drosophila Genetic Reference Panel (DGRP; Mackay et al. 2012; Huang et al. 2014) isogenic lines (Table S1) to produce offspring for use in this experiment. We initially chose these six lines in order to compare three lines exhibiting high cold tolerance in adults but not larvae, and three lines exhibiting high cold tolerance in larvae but not adults (Table S1; Freda et al. 2017). However, initial analyses revealed little evidence for transcriptomic variation tied to variance between these two classes of fly lines, with apparent phenotypic effects highly influenced by outlier lines (Fig. S1). Thus, we treated line (6 levels) as a fixed effect (random effects cannot be modeled using the methods that we applied), providing replication across genetic backgrounds, and did not model phenotypic effects in any of our subsequent analyses.

Each experimental replicate consisted of 10 offspring (10 larvae or 5 male + 5 female adults). To minimize stochastic, environmental effect, each replicate group of 10 offspring was homogenized together to create pools for RNA sequencing. The vial flug (Genesee Scientific, Catalog # 49-102) for each replicate was moistened with water to inhibit desiccation during and after cold exposure. We took an initial sample at 25°C prior to cold exposure (time zero, t0), then exposed all remaining replicates to -5°C by immediately immersing fly vials in a temperature-controlled recirculating bath (ECO RE 2025, Lauda Corporation, Lauda-Königshofen, Germany). We confirmed that vials rapidly reached test temperatures and that larval food did not freeze during treatments (Freda et al., 2017) We then took samples at 30 and 60 minutes during the cold exposure (t30 and t60, respectively). At 60 minutes all remaining vials were removed from the bath and placed back at 25°C, and one final sample was taken 30 minutes after this transfer (30 minutes of recovery, or 90 minutes total, t90). All samples were immediately snap-frozen in liquid N_2_, ground into Tri-reagent (Zymo Research, Irvine CA, USA), then frozen at - 80°C until RNA purification. The overall experimental design included 6 lines by 2 stages by 4 time points by 3 replicates, yielding 144 total samples.

### RNA extraction, library preparation, sequencing, and initial informatics

To extract RNA from DGRP lines for RNASeq, we homogenized each sample (pool of 10 individuals) with a plastic micropestle in Tri-reagent (Zymo) and used the Zymo Direct-zol total RNA extraction kit according to manufacturer’s instructions. We prepared our cDNA libraries using a RNA-tag sequencing approach, as described previously (Lohman et al., 2016). Resulting libraries were sequenced on 5 lanes as 100 bp single-end reads on an Illumina HiSeq 2500 at Kansas University’s Genome Sequencing Core Laboratory, resulting in an average of 6 million reads per sample. The library size for each sample is available in Table S2. We used STAR (Dobin et al., 2013) to map reads to the *D. melanogaster* reference genome (version 6.06) obtained from FlyBase (Gramates et al., 2017), with >95% total mapped reads across all samples. Read counts per gene and per isoform were generated using RSEM (Li and Dewey, 2011). After filtering out all gene models not covered by at least one read in 50% of samples, we retained 13,242 genes. Following normalization of read counts across libraries using the weighted trimmed mean of M-values (TMM) method (Robinson and Oshlack, 2010), we examined variation among libraries using a Multi-Dimensional Scaling (MDS) plot generated using the 500 genes with the highest root-mean-square log_2_-fold change among samples (Robinson et al., 2010). We removed 10 samples that were very clear outliers on the MDS plot (Fig. S2) and exhibited low read counts (less than 200,000 reads) compared to the median read count of 4,808,878 before outliers were removed (Table S2). After removing outliers, all stage × time × line combinations were represented by at least two replicates (Table S3).

### Statistical modelling of temperature- and stage-specific transcription

Our main goal in Experiment 1 was to quantify whether and how the transcriptional response to low temperatures varied between larval and adult life history stages. We expected that many transcripts would be differentially expressed (DE) between life stages because they have very distinct tissue compositions (Arbeitman et al., 2002). Thus, though we estimated gross life stage differences and other contrasts, the parameter of primary interest was a stage × time interaction, indicating stage-specific thermal response during and/or after low temperature exposure (see Fig. 1A for example predicted gene expression trajectories). Below, we detail nested, ad hoc model selection to best estimate that parameter and characterize thermal response trajectories for transcripts with stage-specific expression patterns. The code for this analysis is also publicly available (see Data Availability section). We recognize that transcripts/effects removed from these models may also be of interest, but they were not the focus of this study.

We started with a full generalized linear model with binomial error fitted using the *edgeR* package (Robinson et al., 2010) in R (R Core Team, 2021) to predict the mean read count for each transcript, then removed effects and transcripts to estimate stage × time (interaction) effects that did not depend on DGRP line. The full model included regression coefficients modelling the effects of *stage, time, line*, and all two-way interactions and the three-way interaction of these variables. Statistical inferences from this model identified 130 transcripts with a significant (FDR < 0.05) three-way interaction term. We then removed all transcripts with significant three-way interactions, then fit a reduced model omitting the three-way interaction, which identified 19 transcripts with a significant two-way interaction between *time* and *line*. We removed these transcripts, then fit our final, reduced model including all main effects plus the *stage × time* and *stage × line* two-way interactions.

The transcripts of primary interest in our final, reduced model were those that either 1) had a significant main effect of *time* but no *stage × time* interaction, or 2) had a significant *stage × time* interaction. The former are transcripts that respond to low temperature in similar ways in both life history stages, while the latter are transcripts that exhibit distinct responses to cold in larvae vs. adult flies. We used linear contrasts to estimate the trajectories of differential expression over time for all transcripts in both of these categories by estimating the log_2_ fold change between each time point relative to time zero (t0). This model also allowed us to identify transcripts that had a significant main effect of *stage* or a *stage* × *line* interaction, but no *stage* × *time* interaction. These were not of primary interest, but allowed us to estimate how much of the transcriptome was differentially expressed between life history stages but not responsive to cold. Finally, we used the DAVID functional annotation tool (Huang et al., 2009a; Huang et al., 2009b) to identify functional categories enriched in the set of transcripts illustrating stage-specific responses to cold temperatures.

### Tests for the influence of tissue-specific gene expression

Transcriptomics from whole bodies are coarse measurements that ignore tissue-specificity of gene expression, and in this case differential expression in response to changing temperatures. However, they provide a comprehensive snapshot of whole-organism physiological responses. We could not directly assess how differences in tissue composition across stages might contribute to different transcriptomic responses without tissue-specific RNA libraries. Rather, we tested whether genes that exhibit high levels of tissue-specific expression in *D. melanogaster* were overrepresented in sets of transcripts that we identified as differentially expressed between life stages, or exhibiting stage-by-time interactions.

We quantified tissue specificity of *D. melanogaster* transcripts using data from FlyAtlas2 (Leader et al., 2018) as described in (Cridland et al., 2020). We calculated *τ* for each transcript, a value ranging from 0 to 1, with higher numbers associated with greater tissue-specificity (Yanai et al., 2005). As in (Cridland et al., 2020), if fragments per kilobase of transcript per million mapped reads (FPKM) for whole bodies was less than 2, we set it equal to 2 to avoid inflated estimates for genes with very low expression. We then calculated a normalized expression value for each tissue as the FPKM for that tissue divided by the FPKM for the whole body of the sex/life stage from which the tissue was derived. Finally, we calculated the tissue specificity index, *τ*, as follows:

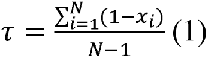

Where *x_i_* is the normalized expression value for the *i^th^* tissue divided by the maximum normalized expression value across tissues and *N* is the number of tissues. We then calculated the median *τ* for a given set of transcripts, e.g., the set exhibiting significant stage-by-time interactions in the above generalized linear models. We compared that point estimate against the median *τ* for 10,000 random samples with the same sample size as the tested set of transcripts to generate a permutation-based p-value.

### Experiment 2: RNAi to test stage-specific functional effects

In order to functionally test whether genes can have stage-specific effects on cold tolerance, we compared the effect of knocking down target gene expression on survival of third instar larvae and 5 day-old adult females and males following a cold stress (Fig 1B). Gene knockdown was achieved using TRiP (Transgenic RNAi Project) lines (Table S1) as described in Teets and Hahn (2018). Briefly, five virgin females from each TRiP line carrying dsRNA under the control of a UAS promoter were crossed to five males of a driver line carrying the GAL4 gene under the control of an actin promoter to produce F_1_ offspring for experiments. The GAL4 driver promotes expression of dsRNA in all tissues to knock down expression of the target gene in the F_1_ generation. We measured survival of groups of 20 larvae or 20 adults (10 adult females and 10 adult males) kept in single fly vials after a 60 min exposure to -5°C, with at least three replicates vials of each stage per line (Fig 1B). The cold treatment was chosen because 40 - 60% of control flies (no RNAi) survived this temperature, allowing us to detect effects of RNAi that either increased or decreased survival relative to the control.

We exposed flies to a -5°C cold stress by immersing vials of larvae and adults in a temperature-controlled Arctic A40 recirculating bath (ThermoFisher, Denver CO, USA) containing 50% (v/v) propylene glycol in water. The fly vials for larvae contained fresh medium, and larvae were allowed to burrow into the food prior to cold treatment via holes poked in the medium; the fly vials for adults were empty (Freda et al., 2017). We verified the temperature in vials using a 36- AWG type-T copper-constantan thermocouple (Omega Engineering, Norwalk CT, USA) interfaced with Picolog v6 software (Pico Technology, Cambridge, UK) via a Pico Technology TC-08 unit. After a 60 min exposure to -5°C, we returned groups of larvae or adult flies to 25°C, 12:12 L:D to recover. Larvae recovered from cold exposure in the same vials and were monitored for adult eclosion over the next 10 d. We classified larval survivors as those that completed development and eclosed as adults (Freda et al., 2017). Adults recovered in small petri dishes containing an approximately 1 cm^3^ piece of fly food medium. We classified adult survivors as those that were motile (could walk/fly independently) 24 h post-cold stress (Jakobs et al., 2015).

### Fly lines

Experiment 2 included 11 TRiP lines (Table S1) whose cold tolerance in adult females has previously been characterized: two control (non-RNAi) lines and nine lines that each knocked down expression of a target gene (Teets and Hahn, 2018). We reasoned that these genes previously had observable effects on adult cold responses, and thus would provide an appropriate test for whether those responses carry over to other life history stages. Two control lines were required because the dsRNA insertion site (and therefore the genetic background) differed among RNAi lines: four lines (+ one control) had an attP2 insertion site, while five lines (+ one control) had an attP40 insertion site (Table S1). Originally, we also planned on knocking out genes identified as having universal or stage-specific responses to cold in Experiment 1, but these experiments were truncated by lab closures during the coronavirus pandemic of 2020.

### Statistical analysis

For each of the nine target genes in Experiment 2 (Table S1), we compared the survival post-cold stress of RNAi and control flies with the same genetic background (attP2 or attP40 insertion sites). We used the *nlme* function in the *lme4* package in R (Bates et al., 2014) to fit generalized linear mixed models with binomial error and a logit link function. We modelled survival as a function of the fixed effects of line (control/RNAi), stage (larvae/adult female/adult male), and their interaction and random (subject level) effects of vial. Example predictions for the effect of RNAi on survival for genes with stage-specific function are in Fig. 1B.

## Results

### Differential gene expression in response to cold is largely stage-specific

A large number of transcripts were significantly (FDR < 0.05) differentially expressed between larval and adult life stages regardless of time sampled during cold treatment (n=10,966, Fig. 2). A smaller, but still sizeable number of transcripts changed in abundance over time. However, only 21 transcripts changed in a similar pattern in both life stages (significant main effect of time, no *stage × time* interaction), while the bulk of the temperature-sensitive transcripts changed over time in a stage-specific manner (n=880 with significant stage × time interaction).

**Figure 2.**
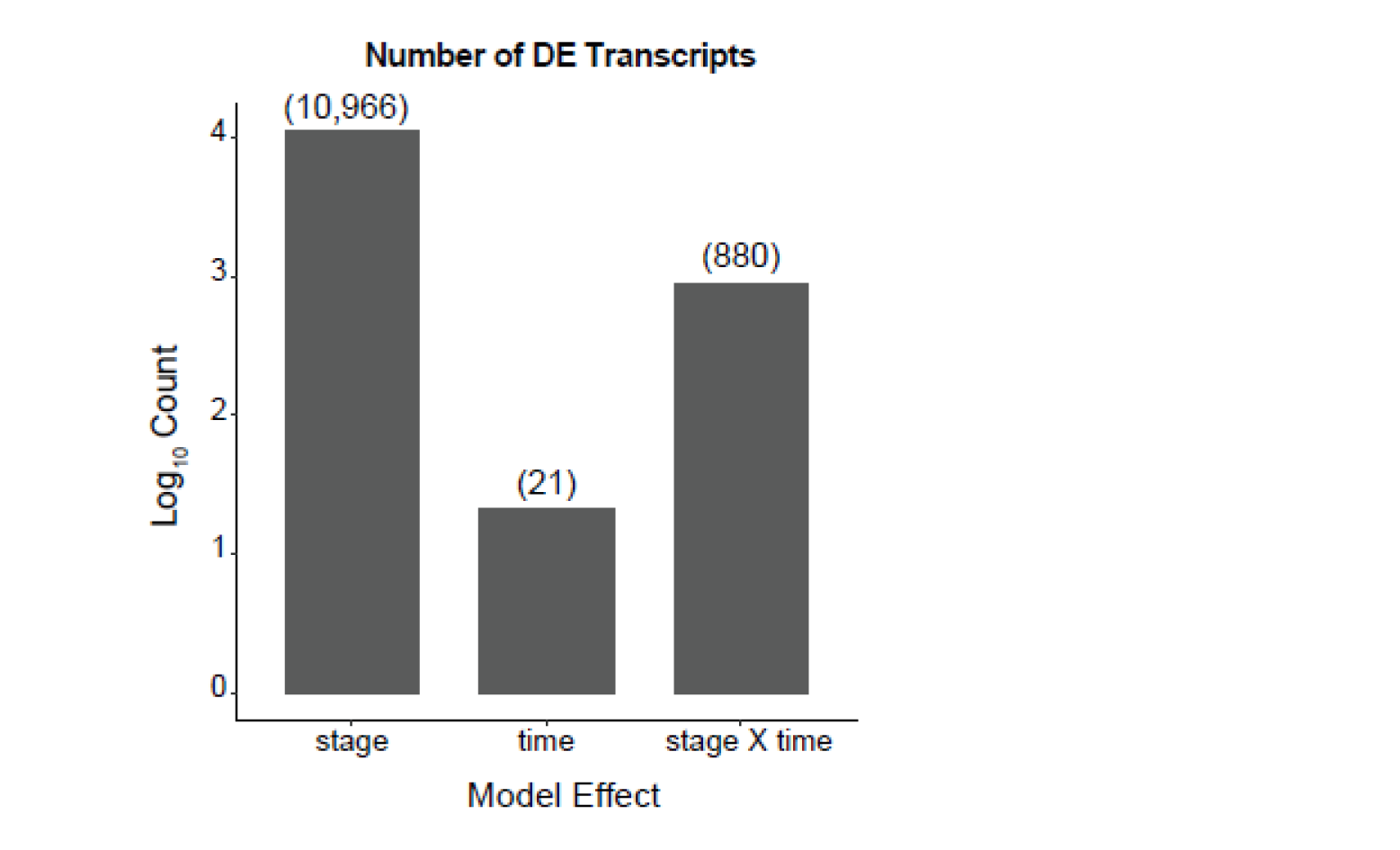
Number of transcripts demonstrating significant (FDR < 0.05) effects of *stage* (larva or adult), *time* (t0, t30, t60, t90), or a *stag*e × *time* interaction during and after cold stress in *D. melanogaster*. The y-axis is log_10_ scaled; actual counts are reported above each bar.

Patterns of change over time were also distinct between life stages. Using linear contrasts, we identified many more transcripts that were significantly DE across at least one time point in larvae (n=763) compared to adults (n=121). Subdividing these into transcripts up-regulated or down-regulated over time on average revealed that most cold-sensitive transcripts in larvae were up- or down-regulated during the cold exposure and remained at similar levels during recovery (Fig. 3A). In contrast, far fewer transcripts were cold-sensitive in adults, and these were mainly up-regulated only during recovery, as has been observed following both cold and heat exposure in other studies of *D. melanogaster* adults (Colinet et al., 2010a; Sinclair et al., 2007; Sørensen et al., 2005; Fig. 3B). Transcripts that changed significantly over time in larvae did not change over time in adults (Fig. 3A; Adult trajectories remain flat). Transcripts significantly up-regulated over time in adults did tend to be up-regulated in larvae as well (Fig. 3B). However, those larval expression trajectories did not demonstrate the same, pronounced up-regulation during recovery observed in adults. The small number of transcripts (n = 21) with significant main effects of *time* but no *stage* × *time* interaction were up-regulated over time in various patterns during cold exposure and recovery (Fig. S3).

**Figure 3.**
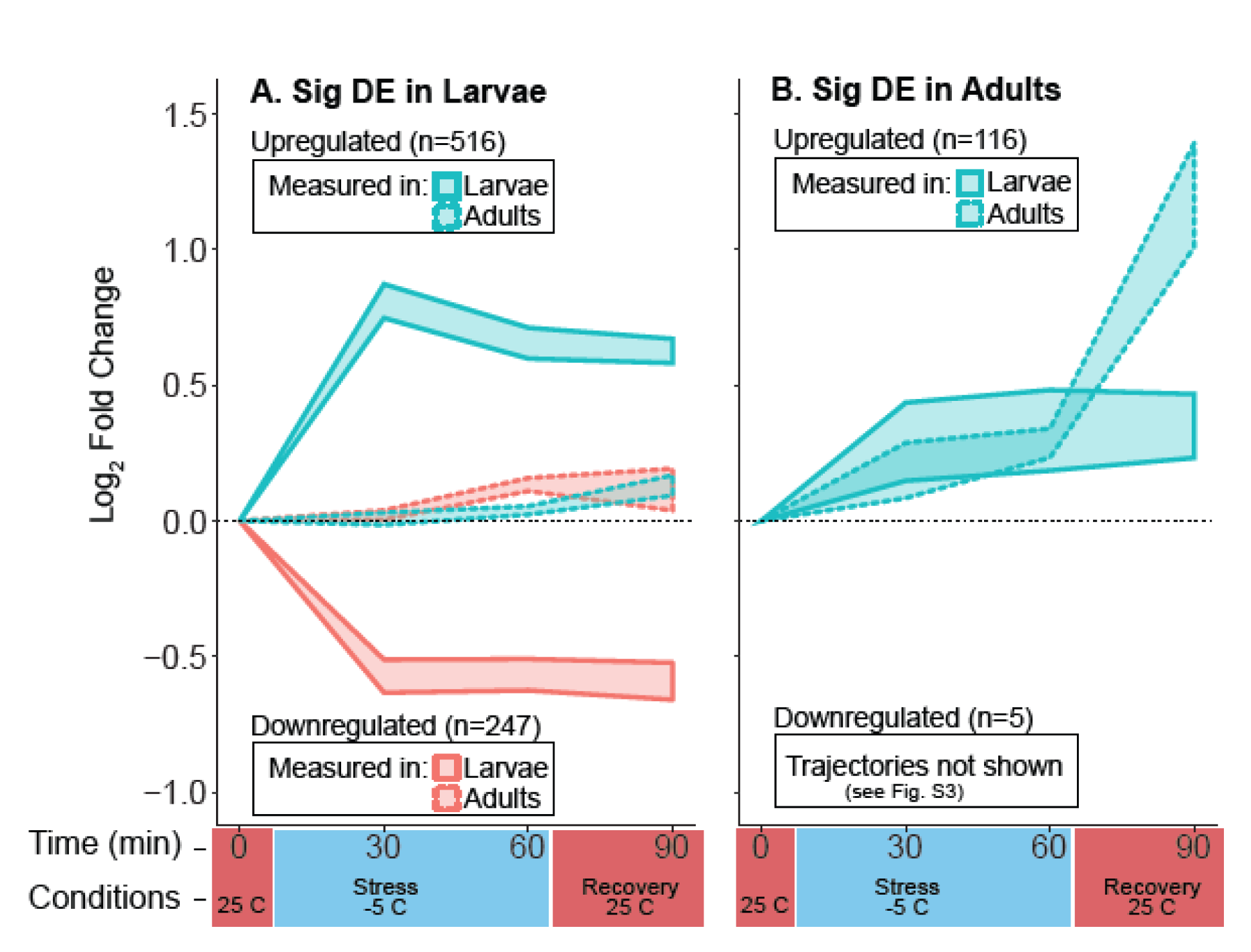
Trajectories of differential expression (DE) during and after cold exposure for the group of transcripts with significant (FDR < 0.05) DE across at least two time points in *D. melanogaster* **(A)** larvae or **(B)** adults. For comparison, the same transcripts measured in larvae (solid borders) and adults (dashed borders) are plotted. Blue indicates transcripts significantly up-regulated on average in (A) larvae or (B) adults, while pink indicates transcripts down- regulated on average in those respective life stages. Y-axis values are log_2_ fold changes at each time point relative to the first (t0) time point. Shaded regions are 95% confidence intervals (c.i.) for each group of transcripts designated in the legends. For example, the blue shaded region with a solid boundary in (A) represents 516 transcripts significantly up-regulated (on average) in larvae, while the blue shaded region with a dashed boundary represents those same transcripts measured in adults. Darker red and blue on the x-axis denote the temperature at which individuals were sampled over the time course.

Closer examination of genes in several functional categories identify candidate mechanisms underlying the cold response that are also stage-specific. Functional enrichment analysis of the 763 and 121 gene significantly differentially expressed over time in larvae and adults, respectively, and with significant *stage × time* interactions identified many overrepresented (FDR < 0.05) functional categories (including Uniprot keyword searches – UPK; Gene Ontology groups – GO; Interpro protein domains – INTERPRO; and Kyoto Encyclopedia of Genes and Genomes pathways – KEGG; Table S4). Below we focus on members of select categories enriched in either larvae (GO Autophagy, INTERPRO Basic leucine zipper, KEGG Fatty Acid metabolism), adults (UPK Stress response), or both (GO Response to bacterium).

Transcripts participating in autophagy, often involved in clearance of cellular damage and nutrient recycling during energy stress (Kroemer et al., 2010), were mainly up-regulated during and after cold exposure in larvae (Fig. 4A). Transcripts with basic leucine zipper domains, largely transcription factors playing roles in developmental regulation, exhibited similar patterns in larvae (Fig. 4B). Transcripts participating in fatty acid metabolism, potentially influencing lipid metabolism or temperature-induced changes in membrane fluidity (Clark and Worland, 2008; Koštál, 2010) were mainly down-regulated during and after cold exposure in larvae (Fig. 4C). All of the transcripts in these three functional categories demonstrated little change during and after cold exposure in adult flies. In contrast, transcripts associated with stress response, mainly chaperonins, exhibited the most pronounced changes only during recovery in adults (Fig. 4D). Though some of these transcripts also changed over time in larvae, many were down- regulated, including multiple copies of the well-known, temperature-inducible stress response gene Hsp70.

**Figure 4.**
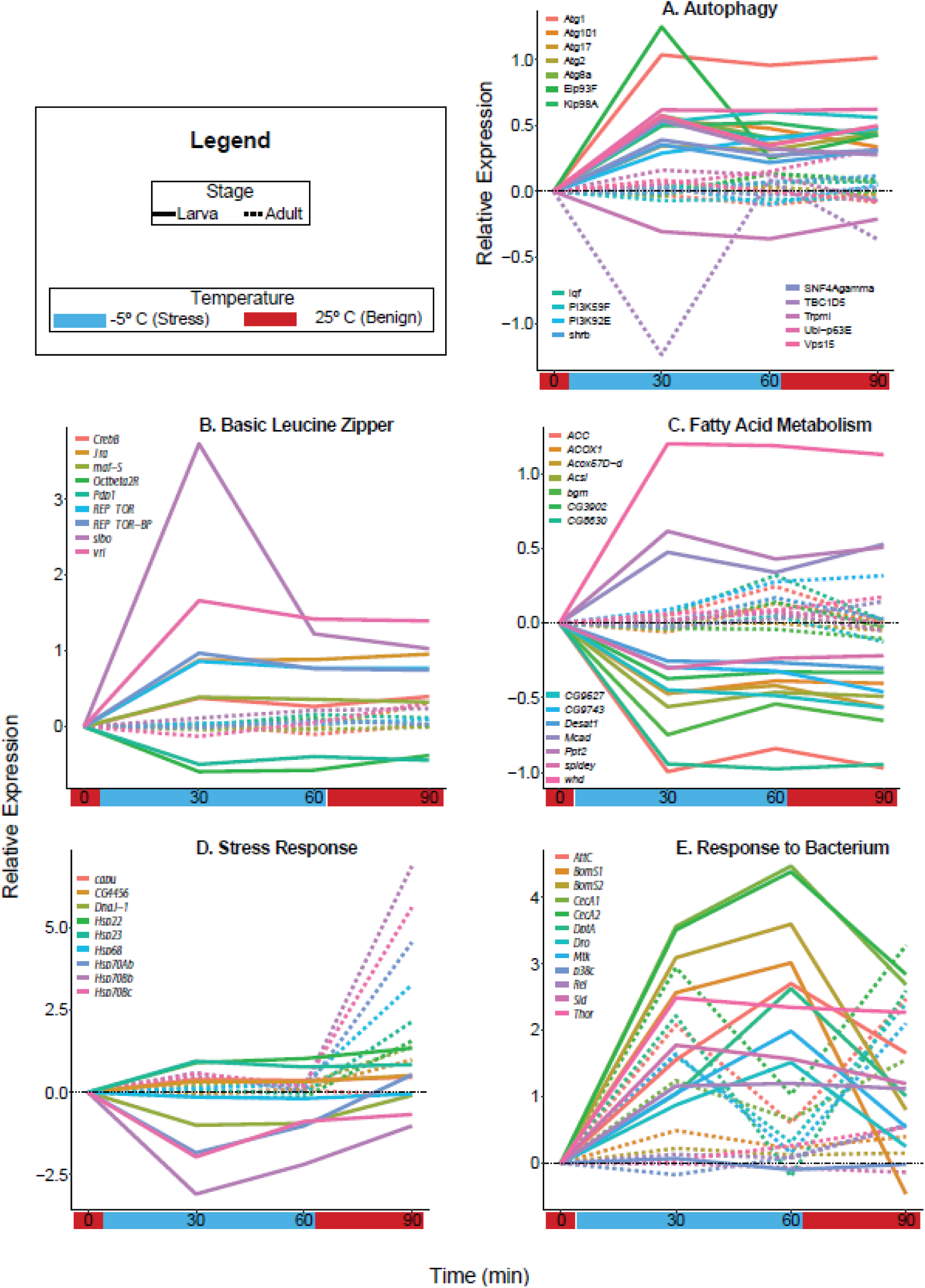
Trajectories of differential expression during (t30, t60) and after (t90) cold exposure for transcripts in select functional categories enriched in the set of all transcripts significantly differentially expressed across at least two time points in *D. melanogaster* larvae or adults. Y-axis values are log_2_ fold changes at each time point relative to the first (t0).

Like transcripts in the stress response category, some transcripts associated with the immune response (within the Response to Bacterium GO group) responded to cold in larvae and adults, though again demonstrating stage-specific patterns (Fig. 4E). The immune response has previously been implicated in responses to thermal extremes in insects (Ferguson et al., 2018; Salehipour-shirazi et al., 2017; Sinclair et al., 2013). All but one transcript in the category were substantially up-regulated during cold exposure in larvae, but tended to decrease in relative abundance during recovery. In adults, transcripts for *Attacin-C* (*AttC*), two Cecropins (*CecA1*, and *CecA2*), *Diptericin A* (*DptA*), and *Metchnikowin* (*Mtk*) were up-regulated at 30 minutes during cold exposure, down-regulated by 60 minutes, then up-regulated again during recovery. One additional Cecropin (*CecC*) was up-regulated over time in a similar pattern for larvae versus adults (main effect of *time* but no *stage* × *time* interaction; Fig. S3).

### Differential expression is unrelated to tissue specificity

We found no evidence that transcripts differentially expressed between whole-body extracts from different stages tended to be more tissue specific. Rather, we found a slight tendency for that set of transcripts to exhibit less tissue specificity than chance expectations. The median *τ* for the set of 10,931 transcripts with stage or stage-by-line effects (FDR < 0.05) was 0.88, and we did not observe a value this small in 10,000 random samples of 10,931 transcripts (median *τ* of random samples = 0.90; p < 0.0001).

Similarly, we found no evidence that transcripts with *stage* × *time* interactions (different responses to the temperature treatments across stages) tended to be more tissue specific. The median *τ* for the set of 849 transcripts with *stage* × *time* effects (FDR < 0.05) was 0.82, and we did not observe a value this small in 10,000 random samples of 849 transcripts (median *τ* of random samples = 0.90; p < 0.0001).

### Some knockdowns had stage-specific effects, but none had consistent cross-stage effects

We observed that the effect of RNAi of target genes on cold tolerance could be stage-specific, although this effect was not universal and was complicated by sex. Three genes of the nine genes tested in this study exhibited clear stage-specific effects of RNAi on cold hardiness (Fig. 5, Table S5). Knockdown of *CG10505* or *Clk* decreased adult, but not larval, survival relative to control flies after a cold stress, suggesting these two genes are important for adult cold tolerance only.

**Figure 5.**
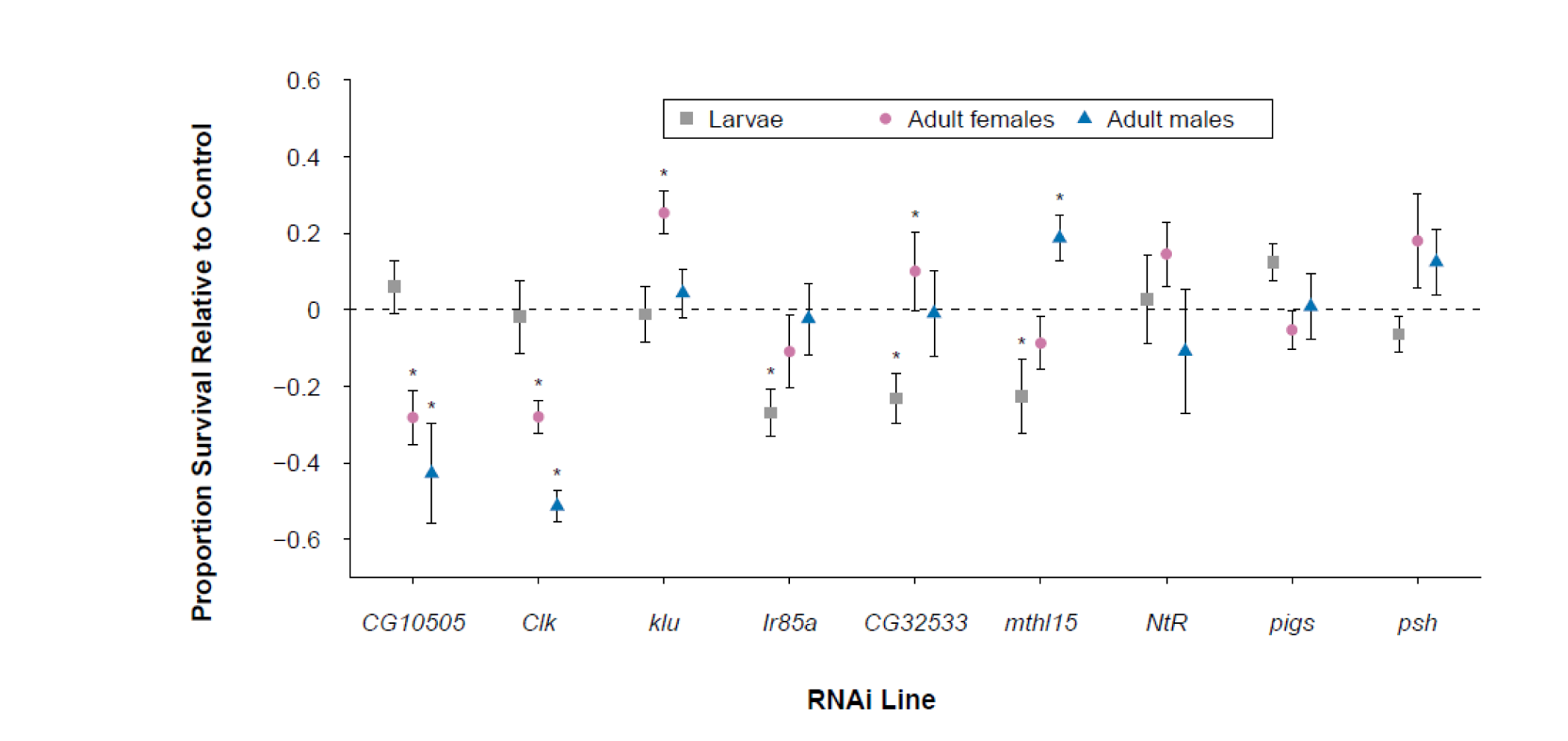
Proportion survival of *D. melanogaster* larvae and adults from lines with RNAi of target gene relative to control lines (no RNAi) following 1 h cold shock at -5°C. Each point represents the mean proportion survival of the RNAi line minus the mean proportion survival of the appropriate control line. Error bars indicate s.e.m., calculated from the proportion survival of three or more replicates of 20 (larvae) or 10 (adult female or male) RNAi individuals. Asterisks indicate a significant effect of RNAi on proportion survival for larvae, adult females, or adult males, based on logistic regressions (Table S4).

Conversely, knockdown of *Ir85a* decreased larval, but not adult, survival, suggesting this gene is important for larval cold tolerance only. RNAi of three other genes had both stage- and sex-specific effects on cold hardiness (Fig. 5, Table S5). *klu* knockdown only increased female adult survival, but had no effect on larvae or male adults. *CG32533* or *mthl15* knockdown had opposite effects on larvae (low survival) and adults (high survival) of one sex only – either female (*CG32533*) or male (*mthl15*). The expression of these two genes therefore seems important for larval cold tolerance but detrimental to either female or male adult cold tolerance. We observed no significant effect of RNAi on cold hardiness for the remaining three genes (*NtR*, *pigs*, *psh*), although *pigs* and *psh* trended toward stage-specific effects (Fig. 5, Table S5).

## Discussion

### Cold tolerance physiology is largely distinct across metamorphosis

Our results generally support the ‘developmentally distinct physiology’ hypothesis, showing that both the expression and function of genes pertinent to cold hardiness differ dramatically across development in *D. melanogaster.* Transcriptional responses to cold in larvae and adults differed in timing (during vs. after cold stress), magnitude (many more DE transcripts in larvae), and constituent genes. In addition, of the nine genes whose expression we knocked down via RNAi, most of them (six) affected adult and larval cold hardiness differently. Though differences in tissue composition across life stages probably have some influence on transcriptional responses to cold, they do not appear to account for the majority of whole-organism transcriptional differences in the thermal response across stages. Other studies have demonstrated transcriptional differences across stages in a complex life cycle (Arbeitman et al., 2002; Chevalier et al., 2006; Sanil et al., 2014; Strode et al., 2006), but this is the first study to our knowledge that demonstrates distinct transcriptome-wide environmental responses across life stages, with additional support from functional genetics experiments.

Although classic CSR genes (e.g. heat shock proteins; HSPs) were not similarly regulated in response to cold in both adults and larvae, we had minor support for the ‘developmentally conserved physiology’ hypothesis based on transcription of immune response genes. Immunity-related genes have been identified as cold-responsive in a number of other studies of adult drosophilid flies (MacMillan et al., 2016; Sinclair et al., 2013; Vermeulen et al., 2013). However, to our knowledge this is the first study to find similar results in adults and larvae, both of which upregulated antimicrobial genes. The function of immunity genes in cold-mediated responses remains unknown, though Vermeulen et al. (2013) suggest that some constituent genes may play a role in repair of cellular damage through their known effects on wound healing. The consistency with which these genes are observed in cold responses across species (Cheng et al., 2017; Salehipour-shirazi et al., 2017; Su et al., 2019; Sun et al., 2019), and here across stages, suggests that they play a specific role in cold physiology, and are not just a general stress response *a la* the CSR.

Though some changes in transcription in response to environmental stress undoubtedly have important, adaptive benefits (Chen et al., 2018; Feder, 1999; Feder and Hofmann, 1999; Feder and Krebs, 1998), differences in baseline (unperturbed) physiology may be equally important. In particular, organisms may have higher fitness when exposed to stress because they are physiologically better prepared prior to stress exposure (Hercus et al., 2003; Krebs and Loeschcke, 1994). To be sure, we have shown that many transcripts differ in expression between stages in benign (baseline) conditions, but this largely reflects the massive developmental differences between the stages. These data do not allow us to identify which of these differences might contribute to differences in expression during and after stress, or to whole organism performance in response to stress, for that matter. However, to the extent that baseline transcriptomes heavily influence transcriptomic responses to a stressor, this still implies that (baseline) physiology affecting cold performance is distinct between life stages.

### Cold hardiness is associated with a muted transcriptional response to cold

Differences between life stages in transcriptomic responses to cold stress likely reflect differences in cold stress resistance between stages. Though not definitively established, a relatively clear pattern is emerging from transcriptomic studies: species or populations that are the most stress resistant are also the least transcriptomically-responsive to environmental stressors. Or, more generally, species or populations that more frequently encounter a given environment tend to have more muted transcriptomic responses to that environment. This is true for Trinidadian guppies responding to predator cues (Ghalambor et al., 2015), fruit-feeding flies responding to different host fruits (Ragland et al., 2015), and marine invertebrates (Lockwood et al., 2010; Schoville et al., 2012), rice plants (Zhang et al., 2012), and other drosophilid flies (Königer and Grath, 2018; Parker et al., 2015) responding to thermal stressors. Adult *D. melanogaster* survive cold stressors better than larvae (Freda et al., 2017; Jensen et al., 2007), and we observed relatively few cold-sensitive transcripts in adults in this study. Moreover, the identity of transcripts involved in the larval transcriptomic response suggest more severe cold-induced damage in larvae compared to adults. Larvae differentially expressed autophagy genes during cold stress, suggesting that larvae need to mitigate cellular damage (i.e. degrade damaged cellular components; Kroemer et al., 2010) or to redistribute macromolecules and energy needed for cell differentiation or growth (Neufeld, 2012; Wang and Levine, 2010). In contrast, adults did not upregulate autophagy-related transcripts and mainly upregulated chaperonins during recovery to preserve cellular function rather than clearing highly damaged cells (Colinet et al., 2010a; Frydenberg et al., 2003; Koštál and Tollarová-Borovanská, 2009). We note that *D. melanogaster* larvae are not susceptible to all stressors; they are more heat-tolerant than adults (Freda et al., 2019), likely because they feed in fruits that can become substantially hotter than air temperatures experienced by adults (Feder et al., 1997). We therefore do not think larvae are more cold-susceptible simply because they are undergoing rapid cellular growth, division, and differentiation compared to adults. Rather, it would appear that each stage has adapted to opposing thermal extremes: heat in larvae and cold in adults.

### Transcriptomic time course and constituent genes differ across life stage

Larvae rapidly differentially regulated a relatively large number of transcripts both during and following cold exposure. These changes likely include active regulation in response to cellular damage, as evidenced by the aforementioned autophagy response. We also observed differential regulation of lipid metabolism in larvae. Fatty acids are important in energy storage (as part of triacylglycerides) and membrane fluidity (as part of phospholipids) (Denlinger and Lee, 2010). Larvae downregulated several desaturases (e.g. *Desat1, CG8630, CG9743*), suggesting that they are not increasing the abundance of unsaturated fatty acids in phospholipids to maintain membrane fluidity at low temperatures (Ohtsu et al., 1998; Overgaard et al., 2005). However, the downregulation of several enzymes associated with fatty acid catabolism (e.g. *ACOX1*) and synthesis (e.g. *ACC, acsl, bgm*) is consistent with restructuring of lipid metabolism to potentially support growth or recovery from stress (Sinclair and Marshall, 2018).

In contrast, adults had relatively muted transcriptomic responses during cold exposure, with a limited (in number of transcripts) but robust (in the degree of differential expression) response during recovery. The best-characterized gene expression response to temperature stress, hot or cold, is upregulation of Hsps and other chaperonins during recovery after exposure to a stressor (Colinet et al., 2010b; Philip and Lee, 2010; Yocum, 2001). This was the most prominent adult response in our study as well, with no detectable changes in Hsp expression during cold exposure. As mentioned above, it is likely that the relative stability of gene expression during stress in adults reflects less severe perturbations from homeostasis and more restricted cellular damage.

### Implications for genetic decoupling across development

Though the transcriptome is only one metric of physiology, the scale of the differences across stages in this study suggests that allelic variants in many genes could strongly affect environmental sensitivity of one stage, while having little effect on other stages. Our results are entirely consistent with empirical studies that repeatedly show little to no genetic correlation in environmental (thermal) sensitivity across metamorphosis in insects (Dierks et al., 2012; Freda et al., 2017; Freda et al., 2019; Gilchrist et al., 1997; Loeschcke and Krebs, 1996; Tucić, 1979). In combination, these results suggest that strong genetic decoupling of environmental sensitivity is relatively common for organisms with complex life cycles, likely facilitating adaptation/acclimation of different life stages to different thermal environments.

Being so widespread, differences in stage-specific thermal tolerance might not appear so surprising. However, temperature is fundamental to limiting species’ spatial distributions (Bale, 2002; Bale et al., 2002), and thus thermal performance must be constrained in some ways.

Though the results of our RNAi knockout experiments suggest that cross-stage pleiotropy for environmental sensitivity is not widespread, we expect that such pleiotropy exists, and will constrain the limits of thermal flexibility across life stages. For example, genetic modifications to increase Hsp70 copy number (and subsequently expression) affected both larval and adult thermal tolerance in *D. melanogaster* (Krebs and Bettencourt, 1999). In that instance, the genetic differences between modified and non-modified lines was relatively extreme (12 extra gene copies). However, there is some evidence for cross-stage effects of naturally segregating genetic variants in plants. Quantitative Trait Locus (QTL) studies in rice have identified QTL associated with cold tolerance at multiple developmental stages, though most QTL only affect a single developmental stage (Yang et al., 2020).

Given the polygenic architecture of environmental tolerance in general (Healy et al., 2018) and thermal tolerance specifically (Barghi et al., 2019; Freda et al., 2017; Sanghera et al., 2011), it’s unlikely that further, detailed analysis of single-locus pleiotropy will fully address questions about the limits of stage-independent adaptations to environmental stressors. Rather, comparative studies leveraging existing variation in stage-specific adaptation or selection studies generating relevant phenotypic variation would seem to be the most promising avenues for further research.

## Acknowledgements

The authors would like to thank Maizey Funk and Lahari Gadey for help with *D. melanogaster* rearing, and the ant-fly-beetle group at CU Denver for comments on earlier versions of the manuscript.

## Data Availability

All raw transcriptomic datasets from this study are available in NCBI BioProject PRJNA783562 (currently embargoed). All code and other data not include in the supplement are available at https://github.com/gjragland/Stage-specific-expression.

## List of symbols and abbreviations

CSR: Cellular Stress Response
DAVID: The Database for Annotation, Visualization and Integrated Discovery
DE: Differentially Expressed
DGRP: Drosophila Genetic Reference Panel
dsRNA: Double-Stranded RNA
FDR: False Discovery Rate
FPKM: Fragments Per Kilobase of transcript per Million mapped reads
GO: Gene Ontology
HSP: Heat Shock Protein
KEGG: Kyoto Encyclopedia of Genes and Genomes
MDS: Multi-Dimensional Scaling
RNAi: RNA Interference
SNP: Single Nucleotide Polymorphism
TRiP: Transgenic RNAi Project

## References

Arbeitman, M. N., Furlong, E. E. M., Imam, F., Johnson, E., Null, B. H., Baker, B. S., Krasnow, M. A., Scott, M. P., Davis, R. W. and White, K. P. (2002). Gene expression during the life cycle of *Drosophila melanogaster*. Science 297, 2270–2275.

Bale, J. S. (2002). Insects and low temperatures: from molecular biology to distributions and abundance. Phil. Trans. R. Soc. Lond., B, Biol. Sci. 357, 849–862.

Bale, J. S., Masters, G. J., Hodkinson, I. D., Awmack, C., Bezemer, T. M., Brown, V. K., Butterfield, J., Buse, A., Coulson, J. C., Farrar, J., et al. (2002). Herbivory in global climate change research: direct effects of rising temperature on insect herbivores. Glob. Change Biol. 8, 1–16.

Barghi, N., Tobler, R., Nolte, V., Jakšić, A. M., Mallard, F., Otte, K. A., Dolezal, M., Taus, T., Kofler, R. and Schlötterer, C. (2019). Genetic redundancy fuels polygenic adaptation in *Drosophila*. PLoS Biol. 17, e3000128.

Bates, D., Mächler, M., Bolker, B. and Walker, S. (2014). Fitting linear mixed-effects models using lme4. arXiv:1406.5823.

Bowler, K. and Terblanche, J. S. (2008). Insect thermal tolerance: what is the role of ontogeny, ageing and senescence? Biol. Rev. 83, 339–355.

Chen, B., Feder, M. E. and Kang, L. (2018). Evolution of heat-shock protein expression underlying adaptive responses to environmental stress. Mol. Ecol. 27, 3040–3054.

Cheng, C.-H., Ye, C.-X., Guo, Z.-X. and Wang, A.-L. (2017). Immune and physiological responses of pufferfish (*Takifugu obscurus*) under cold stress. Fish Shellfish Immunol. 64, 137–145.

Chevalier, S., Martin, A., Leclère, L., Amiel, A. and Houliston, E. (2006). Polarised expression of *FoxB* and *FoxQ2* genes during development of the hydrozoan *Clytia hemisphaerica*. Dev. Genes Evol. 216, 709–720.

Clark, M. S. and Worland, M. R. (2008). How insects survive the cold: molecular mechanisms—a review. J. Comp. Physiol. B 178, 917–933.

Colinet, H., Lee, S. F. and Hoffmann, A. (2010a). Temporal expression of heat shock genes during cold stress and recovery from chill coma in adult *Drosophila melanogaster*. FEBS J. 277, 174–185.

Colinet, H., Lee, S. F. and Hoffmann, A. (2010b). Knocking down expression of *Hsp22* and *Hsp23* by RNA interference affects recovery from chill coma in *Drosophila melanogaster*. J. Exp. Biol. 213, 4146–4150.

Cridland, J. M., Majane, A. C., Sheehy, H. K. and Begun, D. J. (2020). Polymorphism and divergence of novel gene expression patterns in *Drosophila melanogaster*. Genetics 216, 79–93.

Denlinger, D. L. and Lee, J., Richard E. eds. (2010). Low Temperature Biology of Insects. Cambridge: Cambridge University Press.

Dierks, A., Kölzow, N., Franke, K. and Fischer, K. (2012). Does selection on increased cold tolerance in the adult stage confer resistance throughout development? J. Evol. Biol. 25, 1650–1657.

Dobin, A., Davis, C. A., Schlesinger, F., Drenkow, J., Zaleski, C., Jha, S., Batut, P., Chaisson, M. and Gingeras, T. R. (2013). STAR: ultrafast universal RNA-seq aligner. Bioinformatics 29, 15–21.

Feder, M. E. (1999). Organismal, ecological, and evolutionary aspects of Heat-Shock Proteins and the stress response: established conclusions and unresolved issues. Am. Zool. 39, 857–864.

Feder, M. E. and Hofmann, G. E. (1999). Heat-shock proteins, molecular chaperones, and the stress response: evolutionary and ecological physiology. Annu. Rev. Physiol. 61, 243– 282.

Feder, M. E. and Krebs, R. A. (1998). Natural and genetic engineering of the Heat-Shock Protein Hsp70 in *Drosophila melanogaster*: consequences for thermotolerance. Am. Zool. 38, 503–517.

Feder, M. E., Blair, N. and Figueras, H. (1997). Natural thermal stress and heat-shock protein expression in *Drosophila* larvae and pupae. Funct. Ecol. 11, 90–100.

Ferguson, L. V., Kortet, R. and Sinclair, B. J. (2018). Eco-immunology in the cold: the role of immunity in shaping the overwintering survival of ectotherms. J. Exp. Biol. 221, jeb163873.

Freda, P. J., Alex, J. T., Morgan, T. J. and Ragland, G. J. (2017). Genetic decoupling of thermal hardiness across metamorphosis in *Drosophila melanogaster*. Integr. Comp. Biol. 57, 999–1009.

Freda, P. J., Ali, Z. M., Heter, N., Ragland, G. J. and Morgan, T. J. (2019). Stage-specific genotype-by-environment interactions for cold and heat hardiness in *Drosophila melanogaster*. Heredity 123, 479–491.

Frydenberg, J., Hoffmann, A. A. and Loeschcke, V. (2003). DNA sequence variation and latitudinal associations in *hsp23*, *hsp26* and *hsp27* from natural populations of *Drosophila melanogaster*. Mol. Ecol. 12, 2025–2032.

Ghalambor, C. K., Hoke, K. L., Ruell, E. W., Fischer, E. K., Reznick, D. N. and Hughes, K. A. (2015). Non-adaptive plasticity potentiates rapid adaptive evolution of gene expression in nature. Nature 525, 372–375.

Gilchrist, G. W., Huey, R. B. and Partridge, L. (1997). Thermal sensitivity of *Drosophila melanogaster*: evolutionary responses of adults and eggs to laboratory natural selection at different temperatures. Physiol. Zool. 70, 403–414.

Gramates, L. S., Marygold, S. J., Santos, G. dos, Urbano, J.-M., Antonazzo, G., Matthews, A. B., Rey, A. J., Tabone, C. J., Crosby, M. A., Emmert, D. B., et al. (2017). FlyBase at 25: looking to the future. Nucleic Acids Res. 45, D663–D671.

Haldane, J. B. S. (1932). The time of action of genes, and its bearing on some evolutionary problems. Am. Nat. 66, 5–24.

Healy, T. M., Brennan, R. S., Whitehead, A. and Schulte, P. M. (2018). Tolerance traits related to climate change resilience are independent and polygenic. Glob. Chang. Biol. 24, 5348–5360.

Hercus, M. J., Loeschcke, V. and Rattan, S. I. S. (2003). Lifespan extension of *Drosophila melanogaster* through hormesis by repeated mild heat stress. Biogerontology 4, 149–156.

Herrig, D. K., Vertacnik, K. L., Kohrs, A. R. and Linnen, C. R. (2021). Support for the adaptive decoupling hypothesis from whole-transcriptome profiles of a hypermetamorphic and sexually dimorphic insect, *Neodiprion lecontei*. Mol. Ecol. 00, 1– 16.

Huang, D. W., Sherman, B. T. and Lempicki, R. A. (2009a). Systematic and integrative analysis of large gene lists using DAVID bioinformatics resources. Nat. Protoc. 4, 44–57.

Huang, D. W., Sherman, B. T. and Lempicki, R. A. (2009b). Bioinformatics enrichment tools: paths toward the comprehensive functional analysis of large gene lists. Nucleic Acids Res. 37, 1–13.

Jakobs, R., Gariepy, T. D. and Sinclair, B. J. (2015). Adult plasticity of cold tolerance in a continental-temperate population of *Drosophila suzukii*. J. Insect Physiol. 79, 1–9.

Jensen, D., Overgaard, J. and Sørensen, J. G. (2007). The influence of developmental stage on cold shock resistance and ability to cold-harden in *Drosophila melanogaster*. J. Insect Physiol. 53, 179–186.

Kingsolver, J. G., Arthur Woods, H., Buckley, L. B., Potter, K. A., MacLean, H. J. and Higgins, J. K. (2011). Complex life cycles and the responses of insects to climate change. Integr. Comp. Biol. 51, 719–732.

Klockmann, M., Günter, F. and Fischer, K. (2017). Heat resistance throughout ontogeny: body size constrains thermal tolerance. Glob. Chang. Biol. 23, 686–696.

Königer, A. and Grath, S. (2018). Transcriptome analysis reveals candidate genes for cold tolerance in *Drosophila ananassae*. Genes 9, 624.

Koštál, V. (2010). Cell structural modifications in insects at low temperatures. In Low temperature biology of insects (ed. Denlinger, D. L.) and Lee, R. J.), pp. 116–140. Cambridge: Cambridge University Press.

Koštál, V. and Tollarová-Borovanská, M. (2009). The 70 kDa Heat Shock Protein assists during the repair of chilling injury in the insect, *Pyrrhocoris apterus*. PLOS ONE 4, e4546.

Krebs, R. A. and Bettencourt, B. R. (1999). Evolution of thermotolerance and variation in the Heat Shock Protein, Hsp701. Am. Zool. 39, 910–919.

Krebs, R. A. and Loeschcke, V. (1994). Effects of exposure to short-term heat stress on fitness components in *Drosophila melanogaster*. J. Evol. Biol. 7, 39–49.

Krebs, R. A. and Loeschcke, V. (1995). Resistance to thermal stress in preadult *Drosophila buzzatii*: variation among populations and changes in relative resistance across life stages. Biol. J. Linn. Soc. 56, 517–531.

Kroemer, G., Mariño, G. and Levine, B. (2010). Autophagy and the integrated stress response. Mol. Cell 40, 280–293.

Kültz, D. (2005). Molecular and evolutionary casis of the cellular stress response. Annu. Rev. Physiol. 67, 225–257.

Leader, D. P., Krause, S. A., Pandit, A., Davies, S. A. and Dow, J. A. T. (2018). FlyAtlas 2: a new version of the *Drosophila melanogaster* expression atlas with RNA-Seq, miRNA- Seq and sex-specific data. Nucleic Acids Res. 46, D809–D815.

Li, B. and Dewey, C. N. (2011). RSEM: accurate transcript quantification from RNA-Seq data with or without a reference genome. BMC Bioinformatics 12, 323.

Lockwood, B. L., Sanders, J. G. and Somero, G. N. (2010). Transcriptomic responses to heat stress in invasive and native blue mussels (genus *Mytilus*): molecular correlates of invasive success. J. Exp. Biol. 213, 3548–3558.

Loeschcke, V. and Krebs, R. A. (1996). Selection for heat-shock resistance in larval and in adult *Drosophila buzzatii*: comparing direct and indirect responses. Evolution 50, 2354– 2359.

Lohman, B. K., Weber, J. N. and Bolnick, D. I. (2016). Evaluation of TagSeq, a reliable low- cost alternative for RNAseq. Mol. Ecol. Resour. 16, 1315–1321.

MacMillan, H. A., Ferguson, L. V., Nicolai, A., Donini, A., Staples, J. F. and Sinclair, B. J. (2015). Parallel ionoregulatory adjustments underlie phenotypic plasticity and evolution of *Drosophila* cold tolerance. J. Exp. Biol. 218, 423–432.

MacMillan, H. A., Knee, J. M., Dennis, A. B., Udaka, H., Marshall, K. E., Merritt, T. J. S. and Sinclair, B. J. (2016). Cold acclimation wholly reorganizes the *Drosophila melanogaster* transcriptome and metabolome. Sci. Rep. 6, 28999.

McGraw, J. B. and Antonovics, J. (1983). Experimental ecology of *Dryas octopetala* ecotypes: I. ecotypic differentiation and life-cycle stages of selection. J. Ecol. 71, 879–897.

Moran, N. A. (1994). Adaptation and constraint in the complex life cycles of animals. Annu. Rev. Ecol. Syst. 25, 573–600.

Neufeld, T. P. (2012). Autophagy and cell growth – the yin and yang of nutrient responses. J. Cell Sci. 125, 2359–2368.

Nilson, T. L., Sinclair, B. J. and Roberts, S. P. (2006). The effects of carbon dioxide anesthesia and anoxia on rapid cold-hardening and chill coma recovery in *Drosophila melanogaster*. J. Insect Physiol. 52, 1027–1033.

Ohtsu, T., Kimura, M. T. and Katagiri, C. (1998). How *Drosophila* species acquire cold tolerance. Eur. J. Biochem. 252, 608–611.

Overgaard, J. and MacMillan, H. A. (2017). The integrative physiology of insect chill tolerance. Annu. Rev. Physiol. 79, 187–208.

Overgaard, J., Sørensen, J. G., Petersen, S. O., Loeschcke, V. and Holmstrup, M. (2005). Changes in membrane lipid composition following rapid cold hardening in *Drosophila melanogaster*. J. Insect Physiol. 51, 1173–1182.

Parker, D. J., Vesala, L., Ritchie, M. G., Laiho, A., Hoikkala, A. and Kankare, M. (2015). How consistent are the transcriptome changes associated with cold acclimation in two species of the *Drosophila virilis* group? Heredity 115, 13–21.

Philip, B. N. and Lee, R. E. (2010). Changes in abundance of aquaporin-like proteins occurs concomitantly with seasonal acquisition of freeze tolerance in the goldenrod gall fly, *Eurosta solidaginis*. J. Insect Physiol. 56, 679–685.

R Core Team (2021). R: A language and environment for statistical computing. R Foundation for Statistical Computing. Vienna, Austria. URL https://www.R-project.org/

Ragland, G. J. and Kingsolver, J. G. (2008). The effect of fluctuating temperatures on ectotherm life-history traits: comparisons among geographic populations of *Wyeomyia smithii*. Evol. Ecol. Res. 10, 29–44.

Ragland, G. J., Almskaar, K., Vertacnik, K. L., Gough, H. M., Feder, J. L., Hahn, D. A. and Schwarz, D. (2015). Differences in performance and transcriptome-wide gene expression associated with *Rhagoletis* (Diptera: Tephritidae) larvae feeding in alternate host fruit environments. Mol. Ecol. 24, 2759–2776.

Robinson, M. D. and Oshlack, A. (2010). A scaling normalization method for differential expression analysis of RNA-seq data. Genome Biol. 11, R25.

Robinson, M. D., McCarthy, D. J. and Smyth, G. K. (2010). edgeR: a Bioconductor package for differential expression analysis of digital gene expression data. Bioinformatics 26, 139–140.

Salehipour-shirazi, G., Ferguson, L. V. and Sinclair, B. J. (2017). Does cold activate the *Drosophila melanogaster* immune system? J. Insect Physiol. 96, 29–34.

Sanghera, G. S., Wani, S. H., Hussain, W. and Singh, N. B. (2011). Engineering cold stress tolerance in crop plants. Curr. Genomics 12, 30–43.

Sanil, D., Shetty, V. and Shetty, N. J. (2014). Differential expression of glutathione s- transferase enzyme in different life stages of various insecticide-resistant strains of *Anopheles stephensi*: a malaria vector. J. Vector Borne Dis. 9.

Schluter, D., Price, T. D., Rowe, L. and Grant, P. R. (1991). Conflicting selection pressures and life history trade-offs. Proc. R. Soc. Lond. B. 246, 11–17.

Schoville, S. D., Barreto, F. S., Moy, G. W., Wolff, A. and Burton, R. S. (2012). Investigating the molecular basis of local adaptation to thermal stress: population differences in gene expression across the transcriptome of the copepod *Tigriopus californicus*. BMC Evol. Biol. 12, 170.

Sinclair, B. J. and Marshall, K. E. (2018). The many roles of fats in overwintering insects. J. Exp. Biol. 221,.

Sinclair, B. J., Gibbs, A. G. and Roberts, S. P. (2007). Gene transcription during exposure to, and recovery from, cold and desiccation stress in *Drosophila melanogaster*. Insect Mol. Biol. 16, 435–443.

Sinclair, B. J., Ferguson, L. V., Salehipour-shirazi, G. and MacMillan, H. A. (2013). Cross- tolerance and cross-talk in the cold: relating low temperatures to desiccation and immune stress in insects. Integr. Comp. Biol. 53, 545–556.

Sørensen, J. G., Nielsen, M. M., Kruhøffer, M., Justesen, J. and Loeschcke, V. (2005). Full genome gene expression analysis of the heat stress response in *Drosophila melanogaster*. Cell Stress Chaperones 10, 312–328.

Strode, C., Steen, K., Ortelli, F. and Ranson, H. (2006). Differential expression of the detoxification genes in the different life stages of the malaria vector *Anopheles gambiae*. Insect Mol. Biol. 15, 523–530.

Su, Y., Wei, H., Bi, Y., Wang, Y., Zhao, P., Zhang, R., Li, X., Li, J. and Bao, J. (2019). Pre- cold acclimation improves the immune function of trachea and resistance to cold stress in broilers. J. Cell. Physiol. 234, 7198–7212.

Sun, Z., Tan, X., Liu, Q., Ye, H., Zou, C., Xu, M., Zhang, Y. and Ye, C. (2019). Physiological, immune responses and liver lipid metabolism of orange-spotted grouper (*Epinephelus coioides*) under cold stress. Aquaculture 498, 545–555.

Teets, N. M. and Hahn, D. A. (2018). Genetic variation in the shape of cold-survival curves in a single fly population suggests potential for selection from climate variability. J. Evol. Biol. 31, 543–555.

Tucić, N. (1979). Genetic capacity for adaptation to cold resistance at different developmental stages of *Drosophila melanogaster*. Evolution 33, 350–358.

van Gestel, J., Ackermann, M. and Wagner, A. (2019). Microbial life cycles link global modularity in regulation to mosaic evolution. *Nat*. Ecol. Evol. 3, 1184–1196.

Vermeulen, C. J., Sørensen, P., Gagalova, K. K. and Loeschcke, V. (2013). Transcriptomic analysis of inbreeding depression in cold-sensitive *Drosophila melanogaster* shows upregulation of the immune response. J. Evol. Biol. 26, 1890–1902.

Wang, R. C. and Levine, B. (2010). Autophagy in cellular growth control. FEBS Lett. 584, 1417–1426.

Woods, H. A. (2013). Ontogenetic changes in the body temperature of an insect herbivore. Funct. Ecol. 27, 1322–1331.

Yanai, I., Benjamin, H., Shmoish, M., Chalifa-Caspi, V., Shklar, M., Ophir, R., Bar-Even, A., Horn-Saban, S., Safran, M., Domany, E., et al. (2005). Genome-wide midrange transcription profiles reveal expression level relationships in human tissue specification. Bioinformatics 21, 650–659.

Yang, J., Li, D., Liu, H., Liu, Y., Huang, M., Wang, H., Chen, Z. and Guo, T. (2020). Identification of QTLs involved in cold tolerance during the germination and bud stages of rice (*Oryza sativa* L.) via a high-density genetic map. Breed. Sci. 70, 292–302.

Yocum, G. D. (2001). Differential expression of two *HSP70* transcripts in response to cold shock, thermoperiod, and adult diapause in the Colorado potato beetle. J. Insect Physiol. 47, 1139–1145.

Zhang, T., Zhao, X., Wang, W., Pan, Y., Huang, L., Liu, X., Zong, Y., Zhu, L., Yang, D. and Fu, B. (2012). Comparative transcriptome profiling of chilling stress responsiveness in two contrasting rice genotypes. PLOS ONE 7, e43274.

