## Supplementary Figures for "Transcriptomic and functional genetic evidence for distinct ecophysiological responses across complex life cycle stages"

(Supplementary Tables available in SI_Tables.xlsx)

Freda, Philip J.^1*^, Toxopeus, Jantina^2**^, Dowle, Edwina J.^2†^, Ali, Zainab M.^3^, Heter, Nicholas^3^, Lambert-Collier, Rebecca ^3^, Sower, Isaiah^2^, Tucker, Joseph C.^2^, Morgan, Theodore J.^3‡,^, and Ragland, Gregory J.^2^

^1^ Department of Entomology, Kansas State University, 1603 Old Claflin Place, Manhattan, KS 66506, U.S.A.

^*^ *Present address*: Department of Biostatistics, Epidemiology, & Informatics, The Perelman School of Medicine, University of Pennsylvania, D201 Richards Building, 3700 Hamilton Walk, Philadelphia, PA 19104, U.S.A.

^2^ Department of Integrative Biology, University of Colorado Denver, 1151 Arapahoe St., Denver, CO 80204, U.S.A.

^3^ Division of Biology, Kansas State University, 116 Ackert Hall, Manhattan, KS 66506, U.S.A.

^**^ *Present address*: Department of Biology, St. Francis Xavier University, 2321 Notre Dame Ave, NS B2G 2W5, Canada

^†^ *Present address*: Department of Anatomy, University of Otago, 270 Great King Street, Dunedin 9016, New Zealand

^‡^ *Present address*: National Science Foundation, 2415 Eisenhower Avenue Alexandria, VA 22314 U.S.A.

### Supplementary Figures

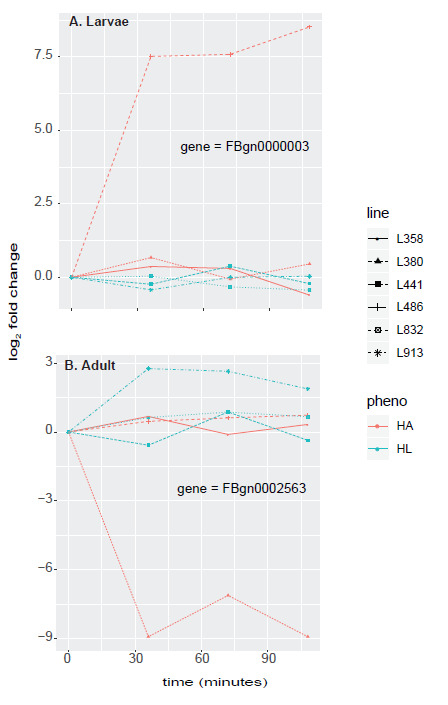

log_2_ fold change

time (minutes)

**Figure S1**. Mean expression trajectories per Drosophila Genetic Reference Panel (DGRP) line, each of which either had high performance as adults (HA) or as larvae (HL). The Y-axis is the log_2_ fold change at each sampling time point compared to the first (0) time point. **(A)** The trajectories for the only gene significantly (FDR < 0.05) differentially expressed between phenotypes from time 0 to time 30 minutes, a time period with active differential expression for larvae (see results in main text). **(B)** The trajectories for one representative gene out of 329 genes significantly differentially expressed between phenotypes from time 0 to time 90 (30 minutes into recovery), a time period with active differential expression for adults (see main text). In both cases, one outlier line is primarily driving differential expression – there are not consistent differences between HA and HL lines in either case. Visual inspection of trajectories for many more genes confirmed that these likely spurious cases of phenotypic effects were common.

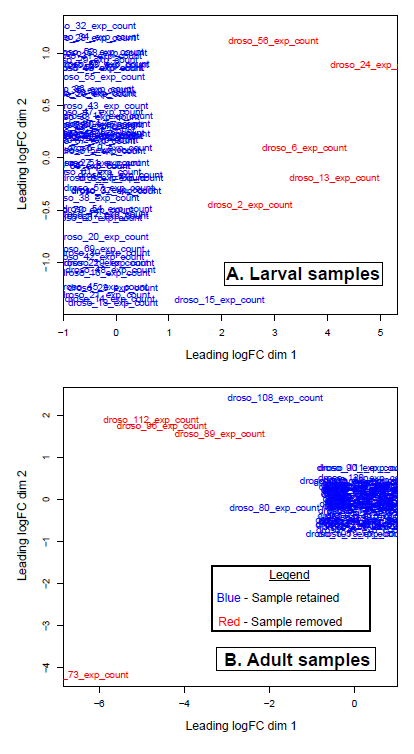

**Figure S2**. Results of Multidimensional Scaling (MDS) analysis of the 500 most differentially expressed transcripts in cold-shocked **(A)** larvae and **(B)** adults. Each RNA library (sample) is plotted as its ID. We removed all samples appearing in red that were clear outliers and had a low (<200,000) number of reads mapping.

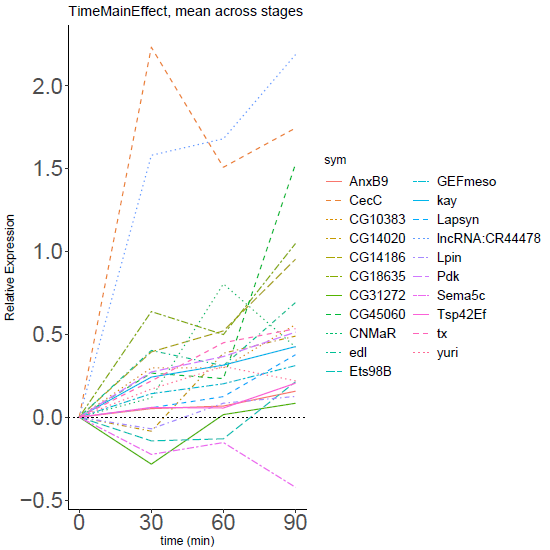

time (minutes)

log_2_ fold change

**Figure S3**. Expression trajectories for each of the 21 genes significantly differentially expressed (FDR < 0.05) during or after cold stress in a similar pattern in larvae and adults (no significant *stage × time* interaction). The Y-axis is the log_2_ fold change at each sampling time point compared to the first (0) time point. Gene abbreviations are consistent with those used in FlyBase.org
